# Investigating risk factors that predict a dog’s fear during veterinary visits

**DOI:** 10.1101/598417

**Authors:** Petra T. Edwards, Susan J. Hazel, Matthew Browne, James Serpell, Michelle L. McArthur, Bradley P. Smith

## Abstract

Attending the veterinary clinic is an integral part of the physical welfare of every companion dog. However, some dogs experience their veterinary visits negatively, which poses a risk of injury to the veterinary staff, their guardian (owner) and themselves during veterinary examinations. It may also influence the regularity of non-urgent veterinary appointments. To date there has been conflicting data on the proportion of dogs that are fearful during their veterinary visits. In this study, we explored the risk factors associated with fear during veterinary examination and in novel situations (including first time at the veterinary clinic) from 26,555 responses in the Canine Behavioral Assessment and Research Questionnaire database. According to their guardians, over half (55%) of companion dogs displayed some form of fearful behaviour (mild-extreme) when examined by a veterinarian, while 14% of dogs exhibited severe or extreme fear. A similar trend was observed with dogs responding fearfully in unfamiliar situations, including the dog’s first time at the veterinary clinic. Chi-squared tests showed every bivariate relationship was significant (p < 0.05). The most important predictors of fear in a veterinary examination were, in order: the dog’s breed group (27.1%), their history of roles or activities (16.7%), where they were sourced (15.2%), their weight (12%), the age of other dogs in the household (9.5%) and dog owner experience (6.3%). However, these risk factors accumulate to explain a total of 7% of variance of fear observed during veterinary examination. Results demonstrate that fear of veterinary visits is common in dogs, but that other factors (including the environment or human-animal interactions) are likely to contribute more to prevalence and severity of this problem than the demographic factors measured here. Finally, we highlight opportunities for future research aimed at facilitating less stressful veterinary visits for dogs and their guardians.

## Introduction

Visits to veterinary clinics are integral to maintaining and improving the health and welfare of domestic dogs. However, the veterinary experience can also be stressful for them. It is currently estimated that between 10% and 78.5% of dogs become stressed or fearful in the veterinary clinic [1–13]. While Doring et al. [4] identified 13% of dogs who refused to enter the veterinary clinic, likely representative of severe fear or distress, Stanford [1] reported that 70% of dogs were unwilling to enter. Further, Mariti et al. [9] observed 29% of dogs displaying ‘extreme’ stress in the waiting room. Such disparity in prevalence is likely a reflection of the methodology employed. For example, variation in the behavioural and physiological measures used to assess stress or fear; the person taking the measurement (e.g., investigator, guardian, veterinary nurse, veterinarian); the locations within the veterinary clinic where stress is measured (e.g., waiting room, examination room or kennels and cages); and the context (e.g. owner present/ absent, mock/real examination). This makes an accurate estimate of the prevalence of stress or fear in dogs visiting veterinary clinics difficult to ascertain.

Negative veterinary experiences can have long-term impacts for the dog, pet owner (guardian) and veterinary staff. A North American study found that the very idea of taking a dog to the veterinary clinic can cause guardians to become stressed (26%). In fact, many guardians (38%) believe that their dog ‘hates’ going to the veterinarian [7]. As such, guardians want to see their veterinarian interact compassionately with their dog [14], especially when certain methods of handling and restraint can be stressful for animals [15–18]. These attitudes and experiences affect guardian decisions about which veterinarian they see and how often they attend [7]. Further, the behavioural or physiological signs of fear and distress can mirror those of pain, illness and some neurological conditions [19], making accurate diagnoses difficult. Not only can stressed or fearful dogs at the veterinary clinic injure themselves, but they also pose a risk of injury to the veterinary staff and their guardians [19–21]. Addressing fear at the veterinary clinic and promoting pet-friendly practice is integral to the continual improvement of companion animal welfare.

With the exception of the dog’s sex [4], size [3], and the benefit of supportive guardian presence [11], our understanding of dog and guardian characteristics that may exacerbate or ameliorate a dog’s fear or stress response at the veterinary clinic is limited. In order to address this shortfall, we explored the proportion of dogs that are fearful during veterinary visits from a large sample of companion dogs. To do this, we analysed two fear-related, veterinary specific questions and their corresponding behavioural subscales from the Canine Behavioral Assessment and Research Questionnaire (C-BARQ). The C-BARQ is a validated research questionnaire available online to dog guardians [20]. It has been used extensively to investigate factors that influence dog behaviour, personality and temperament in general [21–26], as well as to explore how domestication has influenced behaviour more specifically [27]. The C-BARQ has also been used to measure the behavioural effects of neutering in dogs [28, 29], and to investigate the factors associated with aggression [30–32], trainability [33], boldness traits [34], and how training can impact dog intelligence [35], and to explore how dog behaviour or temperament can influence their health and lifespan [36], and the relationship between dogs and their owners [37]. As such, this previously collected, extensive dataset of dogs provides an opportunity to build on our understanding of how dogs experience their veterinary care.

## Method

### C-BARQ

C-BARQ contains 100 items (questions) that have been validated into 14 subscales (factors) of dog behaviour [20]. Guardians respond on a 5-point Likert scale for how serious that behaviour is for their dog or how often it is performed, with ‘0’ being ‘none/ never’ and ‘4’ being ‘extreme/ always’. C-BARQ provides guardians with clear examples of what mild to moderate fear may look like in their dog: “avoiding eye contact, avoidance of the feared object; crouching or cringing with tail lowered or tucked between the legs; whimpering or whining, freezing and shaking or trembling”. Similarly, extreme fear is described as: “exaggerated cowering, and/or vigorous attempts to escape, retreat or hide from the feared object, person or situation”. In this study, C-BARQ responses to two items (questions) related to fear in a veterinary context were analysed to explore the relationship between dog experience at the veterinary clinic and dog and guardian factors (see Table 1 and 2 for details). The items (Questions 43 and 47) asked guardians to report on the extent to which their dog exhibits fearful behaviour during a veterinary examination (Q43; ‘fear of veterinary examination’), and fearful behaviour in unfamiliar situations, including examples of first car trip, first time in elevator, first visit to veterinarian, etc. (Q47; ‘fear of unfamiliar’). The two items, fear of veterinary examination and fear of unfamiliar, loaded onto two different behavioural subscales, touch sensitivity and non-social fear, respectively. Touch sensitivity consists of 4 items (Questions 43, 49, 50 and 51] and refers to dogs that show fearful or wary responses to potentially painful or uncomfortable procedures, including bathing, grooming, nail-clipping and veterinary examinations [20]. Non-social fear contained 6 items (Questions 38, 41, 42, 44, 47 and 48], and refers to dogs that are fearful or wary of sudden or loud noises (e.g. thunder), traffic, and unfamiliar objects and situations [20]. In this study, the predictive value of factors on fear responses in the veterinary context is explored via the two veterinary specific items (Q43 and Q47) in conjunction with the greater insight provided by the two validated behavioural subscales (touch sensitivity and non-social fear).

**Table 1:** Descriptive statistics for mean score of fear (with ‘0’ being ‘no fear’ and ‘4’ being ‘extreme fear’) when examined by a veterinarian (Q.43) and in unfamiliar situations (Q.47) and in the touch sensitivity and non-social fear behavioural subscales according to exposure to variables among 26,555 respondents.

|  | Label | N | % | Fear of veterinary examination | Fear of unfamiliar | Touch sensitivity | Non-social fear |
| --- | --- | --- | --- | --- | --- | --- | --- |
|  |  |  |  | Mean (SD) |  |  |  |
| **Sex** | Male | 13754 | 51.79 | 1.04 (0.010) | 1.01 (0.010) | 0.76 (0.007) | 0.83 (0.007) |
|  | Female | 12801 | 48.21 | 1.09 (0.011) | 1.01 (0.010) | 0.76 (0.007) | 0.86 (0.007) |
| **Age** | Puppy (<0.5yr) | 583 | 2.20 | 0.81 (0.048) | 1.02 (0.046) | 0.67 (0.031) | 0.80 (0.031) |
|  | Adolescent (0.5-3yr) | 12033 | 45.31 | 1.01 (0.011) | 1.05 (0.010) | 0.73 (0.007) | 0.83 (0.007) |
|  | Adult (>3yr) | 13939 | 52.49 | 1.12 (0.010) | 0.97 (0.009) | 0.79 (0.007) | 0.86 (0.007) |
| **Breed group** | Gundogs | 4188 | 15.77 | 0.79 (0.017) | 0.80 (0.015) | 0.58 (0.011) | 0.68 (0.011) |
|  | Hounds | 1481 | 5.58 | 1.16 (0.032) | 1.05 (0.029) | 0.82 (0.021) | 0.86 (0.021) |
|  | Mixed Breed/Unknown | 7370 | 27.75 | 1.33 (0.015) | 1.27 (0.014) | 0.95 (0.010) | 1.04 (0.010) |
|  | Non-Sporting | 2138 | 8.05 | 1.06 (0.026) | 0.99 (0.024) | 0.74 (0.017) | 0.83 (0.016) |
|  | Terrier | 1475 | 5.55 | 0.97 (0.030) | 0.87 (0.028) | 0.74 (0.020) | 0.88 (0.020) |
|  | Toys | 1986 | 7.48 | 1.36 (0.029) | 1.10 (0.026) | 0.97 (0.020) | 0.91 (0.018) |
|  | Utility | 3560 | 13.41 | 0.74 (0.018) | 0.79 (0.017) | 0.57 (0.012) | 0.66 (0.012) |
|  | Working | 4357 | 16.41 | 1.00 (0.018) | 0.95 (0.016) | 0.66 (0.011) | 0.77 (0.011) |
| **Weight** | Smaller (<22kg) | 13331 | 50.20 | 1.21 (0.011) | 1.09 (0.010) | 0.87 (0.007) | 0.93 (0.007) |
|  | Larger (>22kg) | 13224 | 49.80 | 0.91 (0.010) | 0.93 (0.009) | 0.65 (0.006) | 0.75 (0.006) |
| **Neuter status** | No | 6460 | 24.33 | 0.88 (0.014) | 0.88 (0.013) | 0.61 (0.009) | 0.68 (0.009) |
|  | Yes | 20095 | 75.67 | 1.12 (0.009) | 1.05 (0.008) | 0.81 (0.006) | 0.90 (0.006) |
| **Neutered age** | <6 months | 16655 | 62.72 | 1.05 (0.009) | 1.00 (0.009) | 0.75 (0.006) | 0.83 (0.006) |
|  | 6-12 months | 4849 | 18.26 | 1.12 (0.018) | 1.05 (0.016) | 0.80 (0.011) | 0.88 (0.011) |
|  | 12-18 months | 1374 | 5.17 | 1.05 (0.033) | 1.01 (0.030) | 0.75 (0.021) | 0.85 (0.021) |
|  | >18 months | 3677 | 13.85 | 1.07 (0.020) | 0.99 (0.019) | 0.75 (0.013) | 0.85 (0.014) |
| **Reason for neutering** | Birth control | 6864 | 25.85 | 1.10 (0.015) | 1.04 (0.014) | 0.79 (0.010) | 0.89 (0.009) |
|  | Correct behaviour problems | 695 | 2.62 | 1.25 (0.051) | 1.07 (0.045) | 0.87 (0.033) | 0.87 (0.029) |
|  | Correct health problems | 367 | 1.38 | 1.02 (0.063) | 0.85 (0.058) | 0.70 (0.039) | 0.72 (0.040) |
|  | NA | 6809 | 25.64 | 0.89 (0.014) | 0.89 (0.013) | 0.62 (0.009) | 0.69 (0.009) |
|  | Prevent behaviour problems | 796 | 3.00 | 1.16 (0.044) | 1.04 (0.038) | 0.86 (0.029) | 0.87 (0.026) |
|  | Prevent health problems | 1646 | 6.20 | 1.06 (0.030) | 0.90 (0.026) | 0.74 (0.019) | 0.79 (0.017) |
|  | Recommended by veterinarian | 1320 | 4.97 | 1.22 (0.035) | 1.13 (0.032) | 0.88 (0.023) | 0.95 (0.022) |
|  | Required by breeder | 6298 | 23.72 | 1.16 (0.016) | 1.14 (0.015) | 0.84 (0.011) | 0.96 (0.010) |
|  | Unknown | 1439 | 5.42 | 0.98 (0.031) | 0.88 (0.028) | 0.68 (0.020) | 0.80 (0.021) |
| **Source** | Bred by owner | 1083 | 4.08 | 0.73 (0.032) | 0.76 (0.030) | 0.47 (0.020) | 0.56 (0.020) |
|  | Breeder | 10988 | 41.38 | 0.88 (0.011) | 0.83 (0.010) | 0.63 (0.007) | 0.70 (0.007) |
|  | Friend or relative | 2451 | 9.23 | 1.30 (0.026) | 1.16 (0.023) | 0.91 (0.017) | 0.94 (0.016) |
|  | Other | 1420 | 5.35 | 1.05 (0.032) | 1.00 (0.031) | 0.76 (0.021) | 0.85 (0.021) |
|  | Pet store | 922 | 3.47 | 1.37 (0.042) | 1.17 (0.038) | 0.93 (0.028) | 1.05 (0.026) |
|  | Shelter or rescue | 8241 | 31.03 | 1.22 (0.014) | 1.18 (0.013) | 0.89 (0.009) | 1.00 (0.009) |
|  | Stray | 1450 | 5.46 | 1.26 (0.034) | 1.23 (0.031) | 0.92 (0.023) | 1.00 (0.021) |
| **Age when acquired** | Puppy (<0.5yr) | 18842 | 70.95 | 1.03 (0.009) | 0.97 (0.008) | 0.74 (0.006) | 0.81 (0.006) |
|  | Adolescent (0.5-3yr) | 6320 | 23.80 | 1.15 (0.016) | 1.12 (0.015) | 0.82 (0.010) | 0.93 (0.010) |
|  | Adult (>3yr) | 1393 | 5.25 | 1.12 (0.033) | 1.06 (0.032) | 0.80 (0.023) | 0.91 (0.023) |
| **Health problems** | No | 22474 | 84.63 | 1.05 (0.008) | 1.01 (0.007) | 0.75 (0.005) | 0.84 (0.005) |
|  | Yes | 4081 | 15.37 | 1.14 (0.020) | 1.01 (0.018) | 0.80 (0.013) | 0.87 (0.012) |
| **Role** | Breeding & showing | 2247 | 8.46 | 0.65 (0.021) | 0.74 (0.021) | 0.49 (0.014) | 0.59 (0.014) |
|  | Field trials / hunting | 454 | 1.71 | 0.78 (0.050) | 0.72 (0.043) | 0.55 (0.031) | 0.58 (0.030) |
|  | None | 18845 | 70.97 | 1.18 (0.009) | 1.12 (0.008) | 0.85 (0.006) | 0.94 (0.006) |
|  | Other sports | 3545 | 13.35 | 0.92 (0.019) | 0.76 (0.016) | 0.61 (0.011) | 0.65 (0.011) |
|  | Working | 1463 | 5.51 | 0.64 (0.026) | 0.65 (0.024) | 0.49 (0.016) | 0.55 (0.017) |
| **First dog owned** | No | 21728 | 81.82 | 1.01 (0.008) | 0.99 (0.008) | 0.72 (0.005) | 0.82 (0.005) |
|  | Yes | 4827 | 18.18 | 1.30 (0.019) | 1.09 (0.016) | 0.94 (0.012) | 0.95 (0.011) |
| **Owned dog as child** | No | 5281 | 19.89 | 1.12 (0.017) | 1.02 (0.015) | 0.79 (0.011) | 0.87 (0.011) |
|  | Yes | 21274 | 80.11 | 1.05 (0.008) | 1.01 (0.008) | 0.75 (0.005) | 0.84 (0.005) |
| **Age of other household dogs** | NA | 9932 | 37.40 | 1.22 (0.013) | 1.10 (0.011) | 0.87 (0.008) | 0.95 (0.008) |
|  | Older | 5929 | 22.33 | 0.98 (0.015) | 1.05 (0.015) | 0.69 (0.010) | 0.82 (0.010) |
|  | Older and same | 479 | 1.80 | 1.03 (0.056) | 1.07 (0.053) | 0.67 (0.037) | 0.80 (0.040) |
|  | Older and younger | 3001 | 11.30 | 0.86 (0.021) | 0.86 (0.020) | 0.61 (0.014) | 0.68 (0.014) |
|  | Older, younger and same | 748 | 2.82 | 0.83 (0.041) | 0.85 (0.040) | 0.57 (0.028) | 0.63 (0.028) |
|  | Same | 1216 | 4.58 | 1.11 (0.036) | 1.05 (0.033) | 0.78 (0.023) | 0.89 (0.023) |
|  | Younger | 4887 | 18.40 | 1.00 (0.017) | 0.90 (0.015) | 0.74 (0.011) | 0.78 (0.011) |
|  | Younger and same | 363 | 1.37 | 0.99 (0.060) | 0.93 (0.057) | 0.73 (0.043) | 0.76 (0.040) |

**Table 2:**
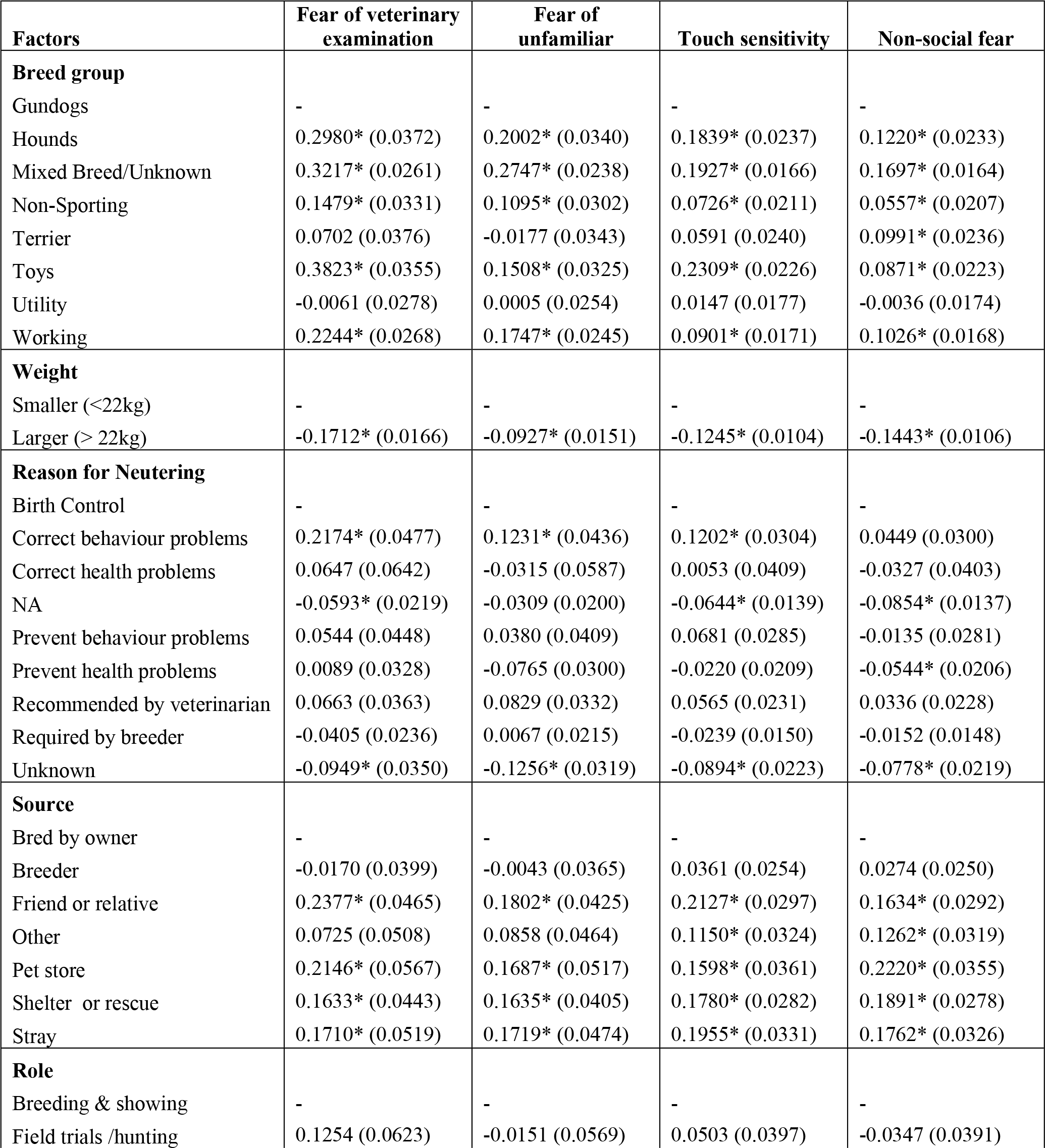
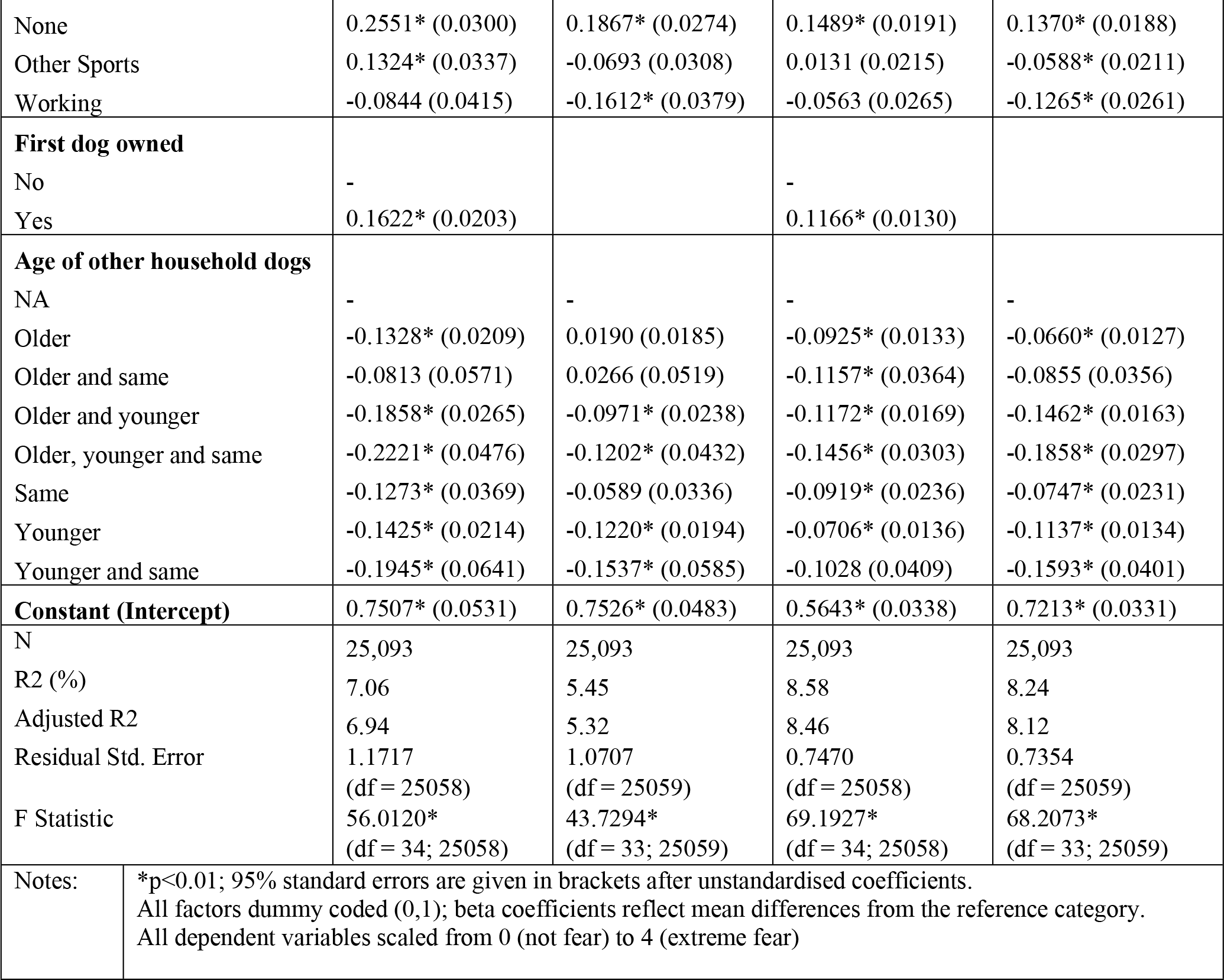
Summary of multivariate regressions predicting dog fear response veterinary examination, unfamiliar situations, touch sensitivity and non-social fear.

### Dog sample

Retrospective data was collected from guardians completing C-BARQ online (https://vetapps.vet.upenn.edu/cbarq/) between 2005 and 2016. Only dog breeds that were clearly identifiable and represented in at least 50 responses were included. Responses that did not include answers to the vital items – Question 43 (fear of veterinary examination) and Question 47 (fear of unfamiliar situations) – were excluded. Breeds were then categorised into breed groups via the Australian National Kennel Council (ANKC) breed list. In the event that a breed was unrecognised by the ANKC, it was categorised in accordance with the American Kennel Club (AKC) (e.g. American Eskimo dog, rat terrier, great Pyrenees, chinook, Spanish water dog). Some breeds were identified in ANKC by a different name (e.g. Belgian sheepdog as groenendael; English bulldog as British bulldog). Breeds not recognised by the ANKC, AKC or the Fédération Cynologique Internationale (FCI) breed lists (English shepherd and American pit bull terrier), were removed. Dog age, neuter age, and age of acquisition were converted into years, and dog weight converted to metric (lbs to kg). C-BARQ is available for guardians with dogs six months and over, and as such, ‘puppies (<6months)’ in this study refer only to dogs that were six months of age at the time of evaluation (N = 583).

### Statistical Analysis

Cross-tabulations were performed between all independent variables and the outcome measures, fear of veterinary examination and fear of unfamiliar. Chi-square tests revealed that every bivariate relationship was significant, due to the large sample size. Accordingly, our analyses focused on determining the magnitude of effect sizes, and assessing the relative importance of each variable in predicting fear of veterinary examination and fear of unfamiliar, in both a univariate and multivariate context. In the first step, we assessed relative importance using two metrics within a framework called *dominance analysis* [38, 39]. The first metric is calculated by entering each variable in isolation (e.g. simple regression), and expressing each model *R*^2^ relatively: e.g. as a percentage of the sum of *R*^2^ explained by all models. The second metric, *lmg* [40], considers the *R*^2^ over all possible combinations of predictors: e.g. given predictors X1, X2, X3, models y ~ X1, y ~ X2, y ~ X3, y ~ X1 + X2, y ~ X2 + X3, y ~ X1 + X3, y ~ X1 + X2+ X3 are considered. Thus, a single predictor’s relative contribution is assessed in terms of the drop in *R*^2^ when it is removed, in all possible multivariate contexts. In the second step, two multiple regression models were fitted for fear of veterinary examination, fear of unfamiliar, touch sensitivity and non-social fear. The first model included all predictors. The second model included only those predictors that contributed more than 5% of the explained variance according to the *lmg* importance metric. Though 5% is an arbitrary threshold for inclusion, in contrast to stepwise methods the *lmg* criteria is a robust variable selection method, because the entire space of possible regression models is evaluated.

## Results

The sample of 26,555 valid responses was evenly distributed by dog sex (51.79% male; Table 1], and the mean dog weight was 22.78kg (median 21.60kg). The mean dog age was 4.52 years (median 4.00 years), mean neuter age was 0.84 years (median 0.48 years) and mean age acquired was 0.77 years (median 0.21 years). The majority of dogs were healthy (84.63%), neutered (75.67%), and purchased as companions (70.97%), with no specific sporting or working role. The most common reasons for neutering were birth control (25.85%), and required by breeder (23.72%). Dogs were most commonly acquired from a breeder (41.38%) and shelter or rescue (31.03%), while the least common source for dogs was a pet store (3.47%), followed by those bred by their guardians (4.08%). The majority of guardians were experienced dog owners, having had dogs previously as adults (81.82%) and/or as children (80.11%). Mixed breeds or dogs of unknown breed were the most commonly reported (27.75%), followed by working breeds (16.41%) and gundogs (15.77%).

Of the 26,555 dogs, over half (55.25%) exhibited some form of fearful behaviour (mild-extreme) when examined by a veterinarian, and 14.23% of guardians reported that their dog showed severe or extreme fear during veterinary examination. Similarly, over half the dogs (57.70%) showed at least some signs of fear in new situations, including potentially the first visit to the veterinary clinic, while 11.02% of dogs displayed severe or extreme fear in unfamiliar situations. In contrast, just over a third of dogs displayed mild-extreme fear for the corresponding subscales of touch sensitivity and non-social fear (35.01% and 37.00% respectively). The mean score of fear for both items (fear of veterinary examination and fear of unfamiliar) and both subscales (touch sensitivity and non-social fear) for each of the independent variables are displayed in Table 1.

### Multivariate regression of fear response

Linear regression models were used to explore the predictive importance of dog and guardian factors in determining fear of veterinary examination and fear of unfamiliar. Table 2 highlights the relationships between the different factors predicting fearful behaviour, analysed through parsimonious regression models for fear of veterinary examination, fear of new situations, touch sensitivity and non-social fear. Considering, ‘all in’ analyses yielded only a slight increase in explained variance of fear responses in comparison to parsimonious models, we focus on the latter. The parsimonious models were significant for all dependent variables, explaining 7.06% of variance in fear of veterinary examination (F= 56.01, df = 34; 25,058, p< 0.01), 8.58% of variance in touch sensitivity subscale (F= 69.19, df = 34; 25,058 p < 0.01), 5.45% of variance of fear of unfamiliar (F = 43.73, df = 33;25,059, p < 0.01), and 8.24% of variance of non-social fear subscale (F = 68.21, df = 33;25,059, p < 0.01). This effect size refers to the proportion of the variation in fearful behaviour that can be attributed to the factors discussed in the following section. For example, approximately 7.06% of the variation of fear observed during veterinary examinations can be attributed to these factors. Likewise, these factors account for 5.45% of the variation in fearful behaviour observed in unfamiliar situations. The intercept represents the grand mean score of fear response for all dependent variables (fear of veterinary examination, fear of unfamiliar, touch sensitivity and non-social fear) for all referents (*B*). The referent score follows the same scale used when guardians reported on each item in C-BARQ, where a score of ‘0’ equates to ‘no fear’, and ‘4’ represents ‘extreme fear’. Other coefficients reflect adjustments to the conditional mean, given each of the predictors.

### Relative importance of factors in explaining variation of fear

The relative importance of each of the factors in predicting fear of veterinary examination, fear of unfamiliar, touch sensitivity and non-social fear are shown in Table 3. While both bivariate (‘first’) and multivariate (‘*lmg*’) analyses are displayed, only the multivariate results are discussed here. Fourteen variables explained more than 5% of the variation in fearful behaviour observed, and are listed in descending order of importance. Only those factors that can be assigned to over 5% of the effect size observed are discussed. A dog’s breed group was the strongest predictor of fear of veterinary examination (27.14%), fear of unfamiliar (26.98%) and touch sensitivity (23.15%). Non-social fear was the only scale in which both role of the dog (24.35%) and dog source (20.02%) explained more of the variance of fear than breed group (18.70%). Role of the dog, dog source, weight and age of other dogs in the household were important factors across all scales. The reason for neutering contributed to the variance of fear observed in all scales, except fear of veterinary examination, while whether the guardian had owned dogs before was only important in fear of veterinary examination and touch sensitivity. Overall, these factors were significant in predicting fear responses in a veterinary context and are important in identifying how dogs experience their veterinary care.

**Table 3:** Relative variable importance in predicting fearful behaviour from C-BARQ items fear of veterinary examination and fear of unfamiliar and subscales touch sensitivity and non-social fear in a bivariate (‘First’) and multivariate (‘*lmg*’) context. *Lmg* scores that indicate the variable captures more than 5% of explained variance in fearfulness are discussed in text.

| Predictor % | Fear of veterinary examination |  | Fear of unfamiliar |  | Touch sensitivity |  | Non-social fear |  |
| --- | --- | --- | --- | --- | --- | --- | --- | --- |
|  | first | *lmg* | first | *lmg* | first | *lmg* | first | *lmg* |
| Breed group | 24.26 | 27.14** | 24.55 | 26.98** | 21.77 | 23.15** | 18.59 | 18.70* |
| Role | 17.36 | 16.68* | 23.34 | 26.81* | 17.40 | 16.79* | 21.16 | 24.35** |
| Source | 16.93 | 15.21* | 21.64 | 19.70* | 17.99 | 17.40* | 20.80 | 20.02* |
| Weight | 10.35 | 11.97* | 4.64 | 5.20* | 10.81 | 14.38* | 7.24 | 10.14* |
| Age of other household dogs | 8.93 | 9.51* | 6.81 | 6.99* | 8.66 | 8.62* | 8.69 | 10.31* |
| Neuter reason | 6.10 | 4.47 | 8.34 | 6.68* | 7.80 | 5.89* | 9.56 | 6.59* |
| Neuter status | 5.18 | 2.28 | 4.60 | 2.25 | 6.58 | 3.53 | 7.85 | 4.56 |
| First dog owned | 6.32 | 6.33* | 1.43 | 1.00 | 6.33 | 6.84* | 2.15 | 1.65 |
| Age when acquired | 1.25 | 0.81 | 3.16 | 2.05 | 1.08 | 0.89 | 2.87 | 1.80 |
| Age | 1.96 | 3.94 | 1.13 | 1.83 | 0.82 | 1.50 | 0.24 | 0.59 |
| Neutered age | 0.34 | 0.31 | 0.31 | 0.39 | 0.31 | 0.48 | 0.36 | 0.43 |
| Health problems | 0.36 | 0.50 | 0.01 | 0.05 | 0.24 | 0.32 | 0.08 | 0.15 |
| Sex | 0.21 | 0.53 | 0.00 | 0.03 | 0.00 | 0.04 | 0.26 | 0.59 |
| Owned dog as a child | 0.46 | 0.32 | 0.05 | 0.03 | 0.22 | 0.16 | 0.15 | 0.12 |
| ** The largest predicting factor |  |  |  |  |  |  |  |  |
| * Factors that contribute over 5% to the variance observed in fear responses to veterinary examination, new situations, non-social fear and touch sensitivity, and are discussed in text |  |  |  |  |  |  |  |  |

### Breed group

Breed group was the largest predictor of fearful behaviour at the veterinary clinic (Table 2). Relative to the other breed groups, toy breeds (*B* = +0.38), mixed breeds (*B* = +0.32) and hounds (*B* = +0.30) predicted the highest scores of fear when examined by a veterinarian. The utility (*B* = −0.01) and gundog (*B* = 0) groups exhibited the least fear during veterinary examination. The same breed group patterns are observed in the corresponding touch sensitivity subscale. However, when assessing fear of unfamiliar situations, mixed/unknown breeds (*B* = +0.28), hounds (*B* = +0.20) and working dogs (*B* = +0.18) displayed the highest scores of fear, while terriers (*B* = −0.02) and gundogs (*B* = 0) exhibited the least fear in new situations. The highest levels of non-social fear were observed in mixed breeds (*B* = +0.17), and hounds (*B* = +0.12), while the lowest non-social fear scores were displayed by utility (*B* = −0.004) and gundogs (*B* = 0).

### A dog’s employment or activity history

The activities or roles a dog has been involved in are the second largest predictor of fear of veterinary examination (16.68%) and fear of new situations (26.81%), and the most important predictor of non-social fear (24.35%). Relative to all roles or activities, dogs used for breeding and showing (*B* = 0) and dogs with a working background (*B* = −0.08) predicted the lowest scores of fear when examined by the veterinarian. Conversely, companion dogs (with no history of formal roles or activities) predicted the highest scores of fear when examined by a veterinarian (*B* = +0.26). Dogs involved in other sports (*B* = +0.13), and field trials or hunting (*B* = +0.13) also tended to exhibit more fear during veterinary examination than working dogs. The same trend was observed in the corresponding touch sensitivity subscale, with companion dogs displaying the highest scores of fear (*B* = +0.15), and those in working roles the least fear (*B* = −0.06). Similarly companion dogs were likely to exhibit the highest fear responses in new situations (*B* = +0.19), and non-social fear (*B* = +0.14), while again, dogs in working roles showed the least fear (*B* = −0.16; *B* = −0.13 respectively).

### Where the dog was acquired

Dogs acquired from a breeder (*B* = −0.02) or bred by their guardians (*B* = 0) predicted the lowest fear scores when examined by a veterinarian. Whereas, dogs acquired from a friend or relative or purchased from a pet store predicted the highest fear scores (*B* = +0.24; *B* = +0.22 respectively). Dogs acquired from a friend or relative were also likely to have higher scores in the touch sensitivity scale (*B* = +0.21), followed by those acquired as a stray (*B* = +0.20), those from a shelter or rescue (*B* = +0.18) and those from a pet store (*B* = +0.16). A slightly different trend is observed in fear of new situations and non-social fear. Dogs acquired from a friend or relative (*B* = +0.18), or as a stray (*B* = +0.17) displayed the highest scores of fear in new situations. In contrast, the highest non-social fear was exhibited by dogs purchased from a pet store (*B* = +0.22), followed by those from a shelter or rescue (*B* = +0.19). Dogs purchased from a breeder were the least fearful of unfamiliar situations (*B* = −0.004), while dogs bred by their guardian had the lowest touch sensitivity and non-social fear scores (*B* = 0).

### Weight

Larger dogs (>22kg) exhibited lower fear scores in comparison to smaller dogs (<22kg) when examined by a veterinarian (*B* = −0.17) and in new situations (*B* = −0.09), and had lower touch sensitivity (*B* = −0.13) and non-social fear (*B* = −0.14).

### Age of other household dogs

Dogs living without conspecifics displayed the most fear across all dependent variables, while dogs that lived with other dogs that were older, younger and the same age showed the lowest scores of fear during veterinary examination (*B* = −0.22) and in the touch sensitivity (*B* = −0.15) and non-social fear (*B* = −0.18) subscales. In contrast, dogs that lived with others that were younger and the same age showed the least fear in new situations (*B* = −0.15).

### Other contributing factors

The reason a dog was neutered also contributed to the variance of fear observed across the majority of variables, but did not contribute over 5% of variance toward fear of veterinary examination. Dogs neutered in order to correct behaviour problems exhibited the highest scores of fear in new situations (*B* = +0.12), touch sensitivity (*B* = +0.12) and non-social fear (*B* = +0.05). Conversely, dogs neutered for unknown reasons displayed the lowest scores of fear in new situations (*B* = −0.13) and in touch sensitivity (*B* = −0.09). Lastly, the guardian’s experience in owning a dog predicts a small proportion of fear observed during veterinary examinations and in touch sensitivity. First time dog owners had dogs that exhibited the highest scores of fear during veterinary examination (*B* = +0.16), and in touch sensitivity (*B* = +0.12), in comparison to guardians that had owned dogs previously.

## Discussion

A large sample size of companion dogs was used to explore the proportion and characteristics of dogs that show a fearful response when visiting a veterinary clinic and in unfamiliar situations. According to their guardians, over half (55%) of companion dogs showed some level of fear when examined by the veterinarian, and 14% exhibited severe or extreme fear in the same context. Likewise, 57% of companion dogs show some form of fear in new situations, including the first time at the veterinary clinic, while 11% exhibit severe-extreme fear. These figures fall within the broad estimates of previous cross-sectional studies [1, 2, 4, 5, 8–11, 13] and arguably provide a more realistic rate of global prevalence of fear of the veterinarian in dogs. In contrast, the touch sensitivity and non-social fear subscales demonstrated a smaller proportion of companion dogs exhibiting some form of fear response (35% and 37% respectively].

The individual items likely measure a wide range of fearful behaviours in dogs visiting a veterinarian as they correlated with different behavioural subscales (touch sensitivity and non-social fear). Fear of veterinary examination likely reflects the association made with handling and potentially painful experience in a clinical setting, while fear of unfamiliar situations (including first time at the veterinary clinic) could reflect a generalised neophobic response. However, the reduced prevalence of fear in the subscales (in comparison to the two individual items), indicates the more general nature of touch sensitivity and non-social fear. That is, while the subscales include items referring to a dog’s veterinary experience, the scales also contain other items that do not. As such, the higher prevalence of fear observed in the individual items reflects the many factors within the veterinary context that may be the cause or catalyst of that fear.

The prevalence of fear in a veterinary context may be influenced by a dog’s genetic predisposition to fear [41]. Godbout et al. [3] identified a small proportion of puppies (10%) that displayed extreme avoidance behaviours during a mock examination. They suggest this likely reflects a proportion of dogs that exhibit anxious behaviours through to adulthood, as a result of a genetic predisposition to an anxious temperament. This is an important area for future research and our results indicate a similar proportion of dogs with severe fear.

As a group, all predictors explained between 5 and 7% of variance in fear of unfamiliar and fear of veterinary examination. The most important predictors were, in order, the dog’s ANKC breed group, the dog’s employment or activity history, where they were sourced, their weight, the age of other dogs in the household, reason for neutering and guardian’s level of experience of dog ownership. As such, each risk factor accumulates to set the foundation of a dog’s predisposition to fearful experience in the veterinary clinic. This low effect size (e.g. 7% for fear of veterinary examination) also highlights the integral role other factors (e.g. environmental, previous experience or human-animal interactions) play in determining the severity of the fear response. This mirrors findings reported in previous studies investigating neuter age and stranger-directed aggression [28] or fear-related behaviours [29], source of acquisition and non-social or stranger-directed fear [42], and litter size and personality [43]. Indeed, Casey et al. [44] suggest the factors associated with human-directed aggression explain a similar amount of variance (<10%), and emphasise that individual experience is likely of much greater importance in determining behaviour.

A dog’s breed group was the best predictor across all the dependent variables. Toy dogs, hounds and mixed breeds were most likely to have the highest scores of fear during veterinary examination in comparison to other breed groups, whereas mixed breeds and hounds predicted higher fear of unfamiliar situations. The same trend was observed in touch sensitivity and non-social fear. Further, according to their guardians, mixed breeds were generally more likely to be fearful of noises in comparison to other breed groups [45]. In contrast, in a cross sectional study, dogs in utility and hound groups were more aggressive to family members than mixed breeds [44]. Although this may simply highlight the difference between aggression and fear, it suggests context is important in determining behaviour – the same dog may react fearfully to an unexpected noise but aggressively toward an unfamiliar person. Additionally, some individual breeds (e.g. dachshunds, Chihuahuas or Jack Russell terriers) have been associated with an increased likelihood of showing aggression toward their guardians and strangers [30]. Considering a dog’s ANKC breed group explained the largest proportion of variance of fear observed, providing additional time within the veterinary consult or extra support to guardians of specific breeds may be valuable in reducing fear in the veterinary clinic.

In the present study, dogs previously employed in working, breeding and showing roles had lower fear of the veterinarian and fear of unfamiliar, while companion animals were most likely to show high levels of fear in the same contexts. The same pattern was observed in touch sensitivity and non-social fear. This is supported by dogs in these roles also having the lowest touch sensitivity scores. The roles that dogs are employed in can influence aspects of their personality and behaviour. Indeed, Lofgren et al. [25] suggest that Labrador retrievers purchased as companions or employed in a gundog role showed higher human and object fear than Labradors that were show dogs, while companion Labradors exhibited greater noise fear than those that were gundogs or show dogs. The reduced risk of fear of veterinary exam and unfamiliar in dogs employed in breeding or working roles may reflect an increased familiarity with procedures associated with veterinary care (i.e. grooming, handling or restraint). As such, we suggest that appropriate handling and grooming practice for companion dogs is equally as important as basic manners training and socialisation in reducing fear in the veterinary context.

The source of acquisition of the dog was also a predictor of fear response in a veterinary context. Dogs acquired from friends or relatives, pet stores shelters and rescues, or as strays were most likely to be fearful during veterinary visits or have high touch sensitivity and non-social fear. In contrast, those bred by their guardian exhibited less fearful behaviour. This reflects a similar finding by Blackwell, Bradshaw and Casey [45], that dogs bred by their guardians less likely to show fear responses to noises than dogs from other sources. Further, puppies from pet stores had increased risk of behavioural issues in comparison to puppies purchased from breeders [42, 46]. Indeed the quality of maternal care can have long term behavioural fallout and alter the physiological responses to stress [47]. Therefore, the puppy’s experience for the first several weeks of life requires careful consideration when investigating the causes of fear during veterinary visits.

Dog size also influences the overall variation of fear observed during veterinary examinations, with lighter (and therefore generally smaller) dogs (<22kg) predicting higher fear scores than heavier dogs (>22kg). Smaller dogs have also been found to be more vocal in the veterinary clinic than larger dogs [3], and are associated with aggressive and excitable, and anxious and fearful behaviour in comparison to larger dogs [48]. However, we highlight that while both breed group and dog weight were predictors of fearful behaviour, the statistical models estimated the effect for breed when controlling for size and vice versa. As such, the extent to which each of the factors contribute to fear of veterinary examination and fear of unfamiliar individually is unknown, and likely inseparable, considering artificial selection for breed phenotype includes size.

The ages of other dogs in the household also contributed to fear responses. Dogs living without conspecifics were most likely to exhibit higher fear in every dependent variable. Lower fear scores for dogs in multi-dog households may reflect social learning. As with the benefit of a positive guardian presence in reducing fear [11], dogs that attend the veterinarian with another familiar, confident dog may take their social cues from that dog and be less fearful. However, whether all dogs in the home attend the veterinarian together, or whether a reduced risk of fear at the veterinary clinic results from some social interaction that occurs at home is unclear. Further, Meyer and Forkman [49] report a link between high social fear in dogs and high guardian emotional closeness with their dog, and so perhaps beneficial guardian or conspecific presence is conditional on the type of attachment. As such, the positive and supportive presence of familiar conspecifics, or a guardian, may help reduce fear during veterinary visits, although further investigation is required.

Guardian level of experience was another contributing factor for fear of veterinary examination and touch sensitivity. Guardians that had never owned a dog previously were more likely to have dogs that exhibited higher fear responses. While over 25% of guardians are able to identify obvious signs of stress in dogs [50], Flint et al. [51] suggest a lack of experience in dog behaviour or attendance at dog training classes is associated with guardians being less likely to identify fear correctly. This suggests a potential for fear to be under-reported in dogs with inexperienced guardians. It also constitutes a significant risk to companion animal welfare as accurately recognising fear is essential in reducing fear in the veterinary context [52–56]. Overall, the predictive weight of guardian experience relative to all other demographic factors is small, although it highlights the importance of guardian education focusing on dog training, behaviour and communication.

Guardians that neutered their dog in order to correct behaviour problems had dogs that were more fearful in new situations, and had higher touch sensitivity and non-social fear. This is supported by Lind et al. [13] who found that dogs with guardian-reported behaviour problems were also rated as more stressed during veterinary visits by the guardian, and the (blinded) test leader. However, as the study design is cross-sectional it may be that dogs that already have behavioural problems, and hence are neutered, are then more likely to be touch sensitive and have non-social fear. Further longitudinal studies are necessary to investigate the influence of neutering on dog behaviour.

While the C-BARQ is a validated questionnaire that clearly describes the behaviour of interest, it is still limited by measurement errors, including: conservative reporting (if guardians are predisposed to report on items like problem behaviours optimistically); guardian interpretation of the items or behaviours; guardians not noticing behaviour during previous veterinary visits, and; time since last veterinary visit. Further, the ability of guardians to accurately identify fear in their own dogs is questionable [9, 50, 51]. Indeed, while Flint et al. [51] found training in recognising fear in dogs resulted in guardians being more likely to correctly identify mild/ moderate and high/extreme fear, they observed no corresponding change in reporting on the guardian’s rating of their own dogs. Conversely, while the current study’s sample size is large, it may reflect responses from guardians that actively seek to know more about their dog’s behaviour, and so, may represent responses from those who are more aware of their dog’s fear or body language. Further, we suggest that the proportion of dogs exhibiting fearful behaviour in the context of unfamiliar situations may be over-representative of experience in the veterinary clinic, as guardians may be reporting on fearful behaviour that occurs in unfamiliar circumstances outside of the veterinary clinic. Future research into dog experience in the veterinary context should corroborate C-BARQ responses for dogs who have recently visited a veterinarian with physiological measures of fear or distress and objective observations.

Overall, it is important to emphasise that the proportion of dogs negatively experiencing their veterinary visits is under-represented by C-BARQ respondents. The items (fear of veterinary examination and fear of unfamiliar situations) within C-BARQ explicitly reflect fear responses only, with no corresponding items for aggression. While several aggression items do refer to grooming or handling by an unfamiliar person (Q14, Q21), they do not expressly mention the veterinarian or veterinary clinic and so were not included in analysis in this study. Aggressive behaviour in the veterinary clinic is also a very real risk for dogs distressed during their veterinary care [55]. As such, it is highly likely guardians with dogs that behave aggressively at the veterinary clinic are not represented in the proportion of dogs experiencing distress during veterinary examination or in unfamiliar situations.

## Conclusion

The results from the present study indicate that around half of companion dogs are experiencing some level of fear or stress when receiving veterinary care. Together, these factors predict approximately 7% of the variation of fear observed during veterinary examinations, and 5% of fear of unfamiliar situations. That is, while they play an important role in determining dog experience in the veterinary context, other factors (e.g. environmental set up or human-animal interactions) are much stronger predictors of a dog’s fear in the veterinary clinic. The cause of the fear in dogs visiting veterinary clinics is still yet to be fully explored. For example, is fear a response to a previous negative experience in a clinic, or are veterinary clinics inherently stressful. While this study has demonstrated the magnitude of the problem, and identified some predictive factors further investigation of individual dogs and the veterinary environment are required.

## References

1. Stanford T. Behavior of Dogs Entering a Veterinary Clinic. Applied Animal Ethology. 1981;7:271–9.

2. Vaisanen M, Valros A, Hakaoja E, Raekallio M, Vainio O. Pre-operative stress in dogs - a preliminary investigation of behavior and heart rate variability in healthy hospitalized dogs. Veterinary Anaesthesia and Analgesia. 2005;32(3):158–67.

3. Godbout M, Palestrini C, Beauchamp G, Frank D. Puppy behavior at the veterinary clinic: A pilot study. Journal of Veterinary Behavior. 2007;2:126–35.

4. Doring D, Roscher A, Scheipl F, Kuchenhoff H, Erhard M. Fear-related behaviour of dogs in veterinary practice. The Veterinary Journal. 2009;182(1):38–43.

5. Hernander L. Factors influencing dogs’ stress level in the waiting room at a veterinary clinic. Department of Animal Environment and Health, 2008.

6. Kim Y, Lee J, Abd el-aty A, Hwang S, Lee J, Lee S. Efficacy of dog-appeasing pheromone (DAP) for ameliorating separation-related behavioural signs in hospitalized dogs. Canadian Veterinary Journal. 2010;51:380–4.

7. Volk J, Felsted K, Thomas J, Siren C. Executive summary of the Bayer veterinary care usage study. Journal of American Veterinary Medical Association. 2011;238(10):1275–82.

8. Hekman J, Karas A, Dreschel N. Salivary cortisol concentrations and behavior in a population of healthy dogs hospitalized for elective procedures. Applied Animal Behaviour Science. 2012;141:149–57.

9. Mariti C, Raspanti E, Zilocchi M, Carlone B, Gazzano A. The assessment of dog welfare in the waiting room of a veterinary clinic. Animal Welfare. 2015;24(3):299–305.

10. Bragg R, Bennett J, Cummings A, Quimby J. Evaluation of the effects of hospital visit stress on physiologic variables in dogs. Journal of American Veterinary Medical Association. 2015;246(2):212–5.

11. Csoltova E, Martineau M, Boissy A, Gilbert C. Behavioral and physiological reactions in dogs to a veterinary examination: Owner-dog interactions improve canine well-being. Physiology Behavior. 2017;177:270–81.

12. Engler W, Bain M. Effect of different types of classical music played at a veterinary hospital on dog behavior and owner satisfaction. Journal of the American Veterinary Medical Association. 2017;251(2):195–200.

13. Lind A, Hydbring-Sandberg E, Forkman B, Keeling L. Assessing stress in dogs during a visit to the veterinary clinic: Correlations between dog behavior in standardized tests and assessments by veterinary staff and owners. Journal of Veterinary Behavior. 2017;17:24–31.

14. McArthur M, Fitzgerald J. Companion animal veterinarians’ use of clinical communication skills. Australian Veterinary Journal. 2013;91(9):374–80.

15. Moberg G. Biological Response to Stress: Implications for Animal Welfare. In: Moberg G, Mench J, editors. The biology of animal stress: basic principles and implications for welfare. USA 2000. p. 1–21.

16. Gregory N. Stress. Physiology and Behaviour of Animal Suffering. United Kingdom: Blackwell Publishing; 2004. p. 12–21.

17. Grandin T. Handling methods and facilities to reduce stress on cattle. Veterinary Clinics of North America: Food Animal Practice. 1998;14(2):325–41.

18. Grandin T. Review: Reducing Handling Stress Improves Both Productivity and Welfare. The Professional Animal Scientist. 1998;14(1):1–21.

19. Frank D. Recognizing behavioral signs of pain and disease: a guide for practitioners. Veterinary Clinics of North America: Small Animal Practice. 2014;44(3):507–24.

20. Hsu Y, Serpell J. Development and validation of a questionnaire for measuring behavior and temperament traits in pet dogs. Journal of American Veterinary Medical Association. 2003;223(9):1293–300.

21. McGreevy P, Georgevsky D, Carrasco J, Valenzuela M, Duffy D, Serpell J. Dog behavior co-varies with height, bodyweight and skull shape. PLoS One. 2013;8(12):1–7.

22. Serpell J, Duffy D. Dog Breeds and Their Behavior. In: Horowitz A, editor. Domestic Dog Cognition and Behavior. Belin Heidelberg: Springer-Verlag; 2014. p. 31–57.

23. Asp H, Fikse W, Nilsson K, Strandberg E. Breed differences in everyday behaviour of dogs. Applied Animal Behaviour Science. 2015;169:69–77.

24. Wiener P, Haskell M. Use of questionnaire-based data to assess dog personality. Journal of Veterinary Behavior. 2016;16:81–5.

25. Lofgren S, Wiener P, Blott S, Sanchez-Molano E, Woolliams J, Clements D, et al. Management and personality in Labrador Retriever dogs. Applied Animal Behaviour Science. 2014;156:44–53.

26. Barnard S, Siracusa C, Reisner I, Valsecchi P, Serpell J. Validity of model devices used to assess canine temperament in behavioral tests. Applied Animal Behaviour Science. 2012;138:79–87.

27. Smith B, Browne M, Serpell J. Owner-reported behavioural characteristics of dingoes (Canis dingo) living as companion animals: A comparison to ‘modern’ and ‘ancient’ dog breeds. Applied Animal Behaviour Science. 2017;187:77–84.

28. Farhoody P, Mallawaarachchi I, Tarwater P, Serpell J, Duffy D, Zink C. Aggression toward Familiar People, Strangers, and Conspecifics in Gonadectomized and Intact Dogs. Frontiers in Veterinary Science. 2017;5(18):1–13.

29. McGreevy P, Wilson B, Starling M, Serpell J. Behavioural risks in male dogs with minimal lifetime exposure to gonadal hormones may complicate population-control benefits of desexing. PLoS One. 2018;13(5).

30. Duffy D, Hsu Y, Serpell J. Breed differences in canine aggression. Applied Animal Behaviour Science. 2008;114(3-4):441–60.

31. Hsu Y, Sun L. Factors associated with aggressive responses in pet dogs. Applied Animal Behaviour Science. 2010;123:108–23.

32. van der Borg J, Beerda B, Ooms M, Silveira de Souza A, van Hagen M, Kemp B. Evaluation of behaviour testing for human directed aggression in dogs. Applied Animal Behaviour Science. 2010;128:78–90.

33. Serpell J, Hsu Y. Effects of breed, sex, and neuter status on trainability in dogs. Anthrozoös. 2005;18(3):196–207.

34. Starling M, Branson N, Thomson P, McGreevy P. “Boldness” in the domestic dog differs among breeds and breed groups. Behavioural Processes. 2013;97:53–62.

35. Marshall-Pescini S, Valsecchi P, Petak I, Accorsi P, Previde E. Does training make you smarter? The effects of training on dogs’ performance (*Canis familiaris*) in a problem solving task. Behavioural Processes. 2008;78:449–54.

36. Dreschel N. The effects of fear and anxiety on health and lifespan in pet dogs. Applied Animal Behaviour Science. 2010;125(3-4):157–62.

37. Hoffman C, Chen P, Serpell J, Jacobsen K. Do dog behavioral characteristics predict the quality of the relationship between dogs and their owners? Human-Animal Interaction Bulletin. 2013;1(1):20–37.

38. Azen R, Budescu D. The Dominance Analysis Approach for Comparing Predictors in Multiple Regression. Pscyhological Methods. 2003;8(2):129–48.

39. Budescu D. Dominance Analysis: A New Approach to the Problem of Relative Importance of Predictors in Multiple Regression. Psychological Bulletin. 1993;114(3):543–51.

40. Gromping U. Variable Importance Assessment in REgression: Linear Regression versus Random Forest. The American Statistician. 2009;63(4):308–19.

41. Serpell J, Duffy DL, Jagoe A. Becoming a dog: Early experience and the development of behavior. In: Serpell J, editor. The Domestic Dog: Its Evolution, Behavior and Interactions with People. 2. Cambridge: Cambridge University Press; 2017. p. 93–117.

42. Wauthier L, WIlliams J. Using the Mini C-BARQ to Investigate the Effects of Puppy Farming on Dog Behaviour. Applied Animal Behaviour Science. 2018.

43. Barnard S, Marshall-Pescini S, Pelosi A, Passalacqua C, Prato-Previde E, Valsecchi P. Breed, sex and litter effects in 2-month old puppies’ behaviour in a standardised open-field test. Scientific Reports. 2018;7(1802).

44. Casey R, Loftus B, Bolster C, Richards G, Blackwell E. Human directed aggression in domestic dogs (Canis familiaris): Occurrence in different contexts and risk factors. Applied Animal Behaviour Science. 2014;152:52–63.

45. Blackwell E, Bradshaw J, Casey R. Fear responses to noises in domestic dogs: Prevalence, risk factors and co-occurrence with other fear related behaviour. Applied Animal Behaviour Science. 2013;145:15–25.

46. McMillan F, Serpell J, Duffy D, Masaoud E, Dohoo I. Differences in behavioral characteristics between dogs obtained as puppies from pet stores and those obtained from noncommercial breeders. Journal of American Veterinary Medical Association. 2013;242(10):1359–63.

47. Czerwinski V, Smith B, Hynd P, Hazel S. The influence of maternal care on stress-related behaviors in domestic dogs: What cacn we learn from the rodent literature? Journal of Veterinary Behavior: Clinical Applications and Research. 2016;14.

48. Arhant C, Bubna-Littitz H, Bartels A, Futschik A, Troxler J. Behaviour of smaller and larger dogs: Effects of training methods, inconsistency of owner behaviour and level of engagement in activities with the dog. Applied Animal Behaviour Science. 2010;123:131–42.

49. Meyer I, Forkman B. Dog and owner characteristics affecting the dog-owner relationship. Journal of Veterinary Behavior. 2014;9:143–50.

50. Mariti C, Gazzano A, Moore J, Baragli P, Chelli L, Sighieri C. Perception of dogs’ stress by their owners. Journal of Veterinary Behavior. 2012;7(4):213–9.

51. Flint H, Coe J, Pearl D, Serpell J, Niel L. Effect of training for dog fear identification on dog owner ratings of fear in familiar and unfamiliar dogs. Applied Animal Behaviour Science. 2018;208:66–74.

52. Overall K. Manual of Clinical Behavioral Medicine for Dogs and Cats. USA: Elsevier; 2013.

53. Yin S. Low Stress Handling, Restraint and Behavior Mmodification of Dogs and Cats: Techniques for Developing Patients Who Love Their Visits. Kolus C, Adelman B, editors. USA: CattleDog Publishing; 2009.

54. Shepherd K. Behavioural medicine as an integral part of veerinary practice. In: Horowitz D, Mills D, editors. BSAVA Manual of Canine and Feline Behavioural Medicine. 2nd ed. England: BSAVA; 2009.

55. Moffat K. Addressing canine and feline aggression in the veterinary clinic. Veterinary Clinics of North America: Small Animal Practice. 2008;38(5):983–1003.

56. Lloyd J. Minimising Stress for Patients in the Veterinary Hospital: Why It Is Important and What Can Be Done about It. Veterinary Sciences. 2017;4(2):1–19.

